# Production benefits of community-created vermicompost and worm juice in a South African township

**DOI:** 10.64898/2026.02.05.704121

**Authors:** Adam D. Kay, Wezo Sinqomo, Sithembele Mnguni, Lindokuhle Mpayipeli, Muna Abdullahi, Shaylee Beckfield, Justa Heinen-Kay

## Abstract

As urbanization increases globally, many residents of cities face challenges from food insecurity and general poverty. In sub-Saharan Africa (SSA), many migrants to cities come from rural, agricultural backgrounds and may be eager to farm in community or kitchen gardens for nutritional, economic, and social benefits. However, soil nutrient deficiencies and other constraints on production limit potential gains from urban agriculture. As a result, low-cost innovations to increase production could provide important community benefits. Here, we report on results from an experiment to assess the production value of worm-generated, food-waste compost created by a community initiative in Khayelitsha, South Africa, one of the largest and fastest growing urban settlements in South Africa. We first describe a community-embedded program for generating worm-based compost (“vermicompost”) and “worm juice” (liquid extracts from the composting process). We then describe an experiment conducted in the community in which we field tested the effects of vermicompost and worm juice addition on the growth of Swiss chard, a common winter crop at this location. We grew plants in one of three levels of vermicompost addition (0%, 10%, 20%) either with or without worm juice in a fully factorial design. We found that 10% and 20% vermicompost significantly (P < 0.0001) increased chard yield (by ∼51% and ∼59% respectively), and worm juice also significantly increased yield (p = 0.024). Soil analyses indicated that vermicompost addition reduced soil pH and increased soil P levels. These results suggest that a community-embedded, grassroots vermicomposting operation can substantially increase yields for urban farmers, thereby increasing the economic and social benefits that emerge from community and kitchen gardens in marginalized urban areas.

## Introduction

Rapid urbanization is one of the driving forces of social change in the 21st century. Over half of the global human population currently lives in urban areas, with major consequences for human health and economic opportunity (Elmqvist et al. 2015, McPhearson et al. 2016, Fan et al. 2025) and for the global environment (Kronenberg et al. 2024). Although the primary motivation for much of recent rural-to-urban migration is economic opportunity (Potts 2016, Zimmer et al. 2020), many new migrants and longer established residents struggle in the face of growing economic and social disparities in urban areas. These trends are reflected in the rapid expansion of informal settlements in cities around the world (Marx et al. 2013, Williams et al. 2019), with many of the fastest growing settlements in sub-Saharan Africa (SSA) (Büttner et al. 2025). Creating economic opportunities and improving social services and environmental conditions in such settlements is essential for ensuring social and environmental resiliency in the coming decades (Rigon 2022, G and Sonda 2024).

One proposed tool for improving economic, social, and environmental conditions in SSA urban settlements is urban agriculture (Mwakiwa et al. 2018, Modibedi et al. 2021, Musosa et al. 2022). Urban food insecurity and poverty alleviation are especially pressing challenges in SSA cities, where residents face multiple stressors including limited access to employment and social services, high food prices, and scarcity of land for home gardening (Kiribou et al. 2024). Expansion of urban agriculture in SSA cities has been shown in multiple studies to improve farmer nutrition, generate supplemental household income, and build community connections (Olivier 2019, Swanepoel et al. 2021, Kanosvamhira and Shade 2024, Kanosvamhira 2024). However, multiple challenges limit urban agriculture in marginalized SSA urban areas, including property crime, land tenure insecurity, and constraints on crop production (including high input costs, water scarcity, soil contamination, and soil nutrient deficiencies) (Reuther and Dewar 2006, Olivier 2019, Kanosvamhira 2023). Innovations to reduce or eliminate these challenges could increase material and social benefits of engaging in urban agriculture and attract a new generation of urban farmers (Kanosvamhira 2023).

Here, we report on a field test of products from a low-tech, grassroots program that aims to increase crop production for urban farmers in Khayelitsha, South Africa, one of the largest and fastest growing marginalized urban areas in South Africa (Lynge et al. 2022). Khayelitsha was established on the periphery of Cape Town in 1983 as part of a forced-removal program by the apartheid government of South Africa. Although much of the Cape Town area has been deemed “very suitable” for urban agriculture (Kanosvamhira et al. 2025), many of the forced-removal program settlements were created on areas with sandy soils lacking sufficient organic matter to support productive agriculture (Kanosvamhira 2023). In a recent survey of 100 households in Khayelitsha, only 55% of households that grew crops indicated that they added any fertilizer for their crops, all of those who added fertilizer would like to use more, and all growers indicated that cost was the most important constraint on fertilizer use (Kay et al in preparation).

One potentially scalable opportunity to broadly increase fertility in nutrient-poor, urban soils is through the production and application of compost derived from food waste. Food waste can be significant in SSA cities (Sheahan and Barrett CB 2017, Mmereki et al. 2024) with important sanitary, public health, and environmental consequences (Chalak et al. 2016, Mmereki et al. 2016). Composting urban food waste has the potential to lessen these consequences while providing a substantial amount of valuable product for soil amendment (Cerda et al. 2018, Mmereki et al. 2024).

One promising approach for converting urban food waste into a soil amendment for urban agriculture is through worm-based composting (“vermicomposting”) (Garg et al. 2012, Schröder et al. 2021, Wang et al. 2024). Composting worms (*Eisenia fetida*) can break down a variety of fruit and vegetable scraps, and their waste products (castings) and liquid extracts (here referred to as “worm juice”) are rich in microorganisms, nutrients, and substances promoting plant growth (Ganapathy et al. 2025). Compared to traditional composting, vermicomposting is superior for nutrient recycling because of the worms’ ability to quickly mineralize organic waste, releasing nutrients that plants can absorb (Wang et al. 2024). Additionally, vermicomposting can be conducted without creating food waste odor or attracting pests, making it a viable option for urban environments (Sherman 2018). Vermicomposting is particularly well-suited for the Cape Town region because its moderate climate stays within the temperature tolerance range of the *Eisenia fetida* (red wiggler) worms that are used for vermicomposting (13-27°C, Sherman 2018), thus allowing for year-round outdoor operations. Finally, vermicomposting projects can in theory be created at a micro scale with minimal infrastructure, making them potentially beneficial to marginalized urban communities in sub-Saharan Africa and beyond.

Here, we present results from a field test investigating the effects on crop yield by applying the vermicompost and worm juice produced by a grassroots, community-embedded vermicomposting operation. We include a description of the operation and of how the vermicomposting products were created. We also discuss opportunities for future development and refinement of products to suggest the potential for such a decentralized operation to benefit urban farmers in marginalized urban areas.

## Methods

### Setting

Khayelitsha is one of the largest townships in South Africa. Its population was estimated at ∼400,000 in 2011 (City of Cape Town 2011) but has likely grown considerably since then. Average household income in Khayelitsha is less than 160 rand (∼$10 USD)/day (Lynge et al. 2022) and unemployment is likely between 50 and 70% (Webb 2021).

### Vermicompost and worm juice production

We created vermicompost using a partially decentralized system involving the local community (described further in Kay et al. in preparation). The operation consists of a network of local residents who begin the composting process at their homes, coupled with a central site where composting is finished (Figure 2). This operation was established by two Khayelitsha residents (co-authors Mnguni and Sinqomo) and co-authors Kay and Heinen-Kay in Ward 94 in Khayelitsha. Below we briefly describe the development of the operation and the process by which vermicompost and worm juice were created for this study.

**Figure 1.**
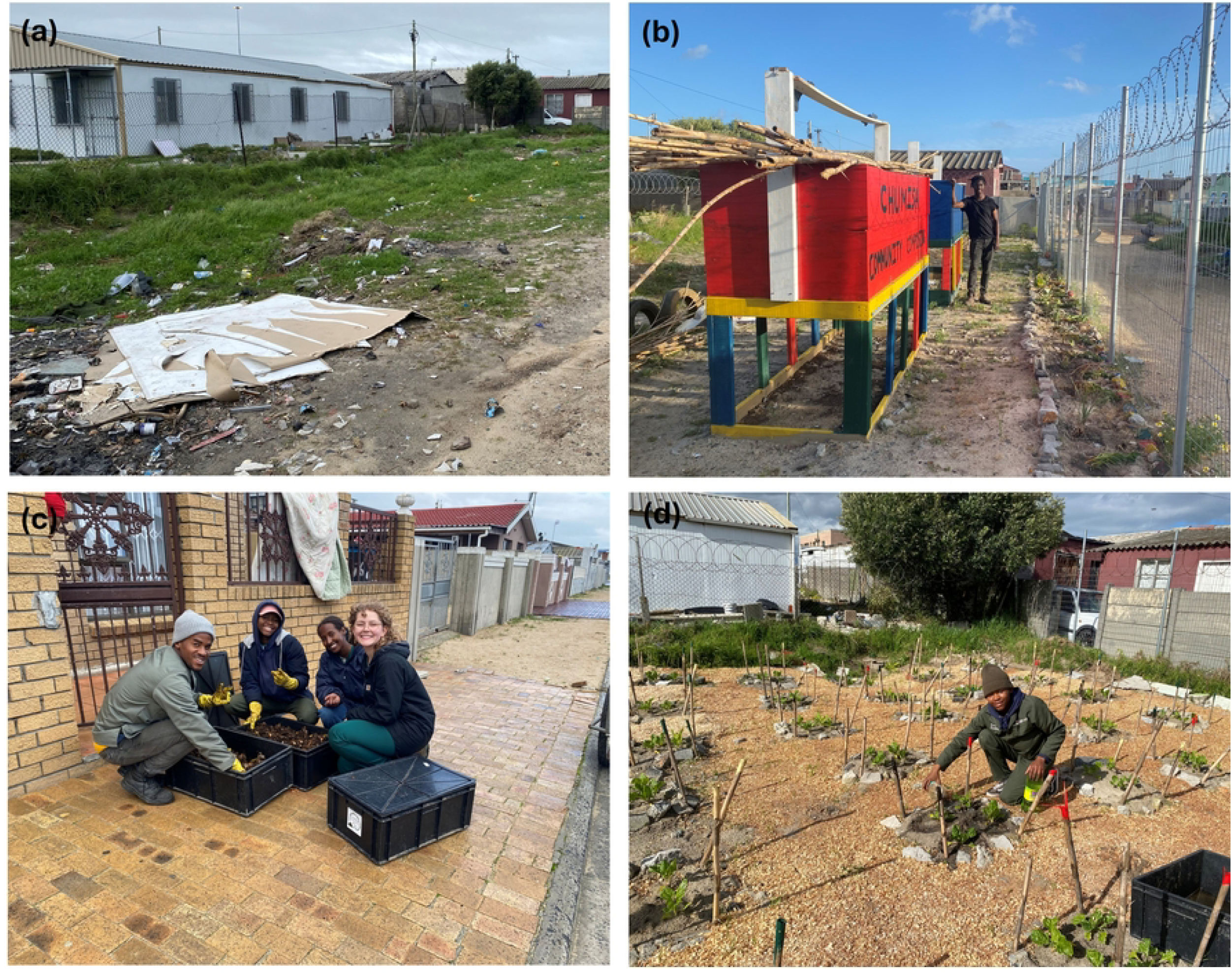
Depictions of the vermicomposting operation in Khayelitsha, South Africa described in this study. A) The lot in Ward 94 in Khayelitsha before establishment of the site for creating finished vermicompost used in this study. B) Picture of the central site with continuous-flow vermicomposting system, with co-author Mnguni. C) A home visit for collecting “pre-compost” with co-authors Sinqomo, Mpayipeli, Abdullahi, and Beckfield. D) The growing site, with co-author Mpayipeli.

**Figure 2.**
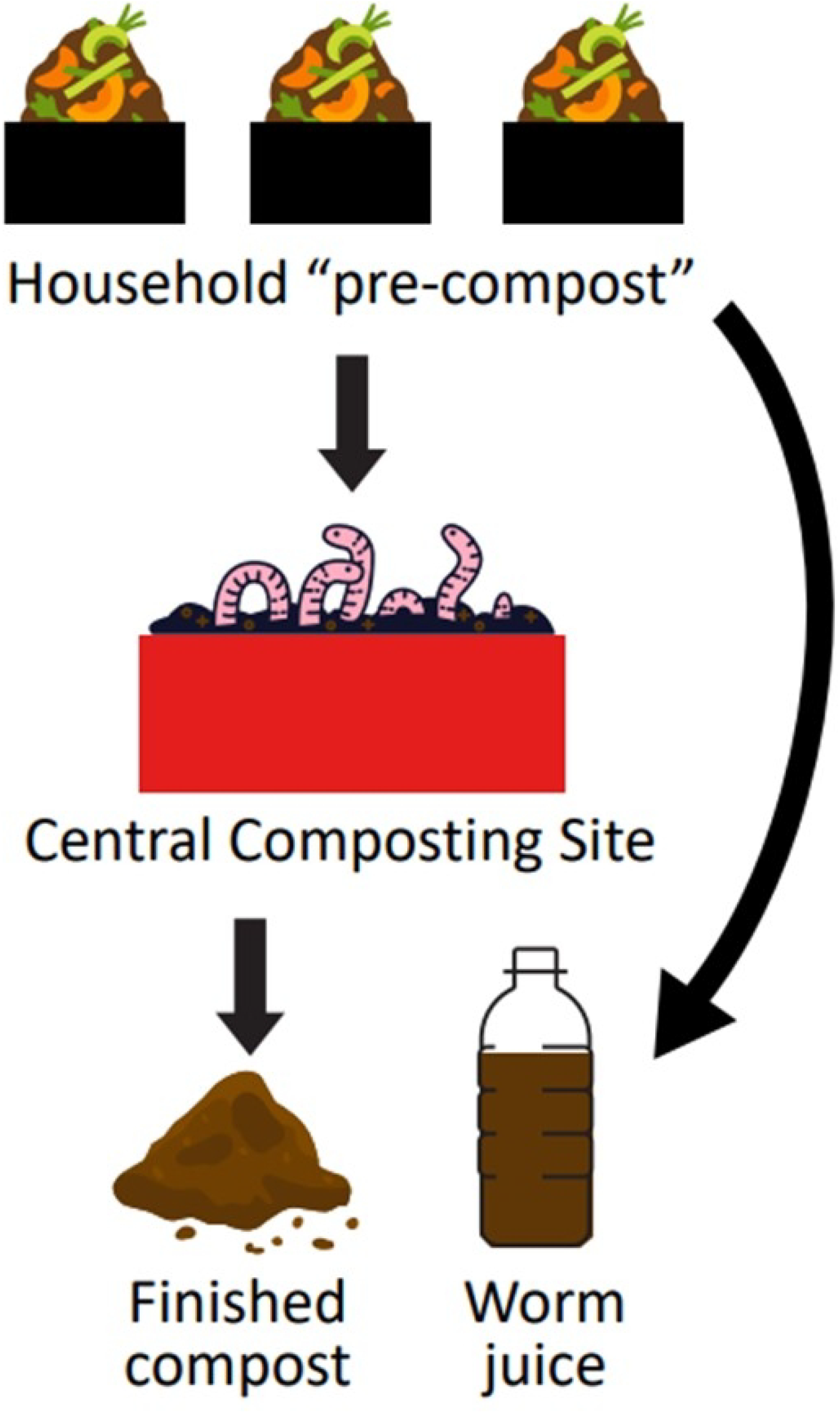
Graphic of the vermicompost and worm juice generation process. About 60 households contribute “pre-compost” (partially worm composted food waste and cardboard) that is transferred to larger continuous-flow vermicomposting systems at a central site. Some households also regularly contributed worm juice (liquid extracts collected from the base of home vermicomposting systems)

We created a central site in August 2023 on a vacant lot previously used as a dumping site for household trash (Figure 1A). After securing the site, we built 2 2m × 1m × 1m continuous-flow vermicomposting systems (Figure 1B) and then added ∼30,000 red wriggler (*E. fetida*) worms, cardboard, bedding, and fruit and vegetable waste collected from local households. We then maintained this system for ∼10 months to build up the worm population using as substrate food waste from local households and cardboard scraps discarded in surrounding areas.

We then added local households to the composting process starting in June 2024. We asked households if they would be interested in composting their food waste at home if we provided them with all supplies and helped them manage their operation. We first recruited 6 elders from the community to participate. We supplied each household with two 82 × 46 × 30cm plastic bins, all necessary supplies (coco coir bedding, cardboard pieces, and ∼500 worms) for composting, and basic instructions. After, we visited households each week to provide any needed supplies (i.e., bedding, cardboard, or extra worms) and to retrieve 30-50% of accumulated “pre-compost” (a mixture of new food waste and previously composted materials) (Figure 1C), which was then returned to the central site and added to the continuous-flow systems. Between June 2024 and May 2025, we expanded the program to include 60 households within walking distance of the central site, with most of these households contributing “pre-compost” each week.

Here, we report on a field test of the yield value of vermicompost products (finished vermicompost, worm juice) created from this operation. In June 2025, we harvested finished vermicompost from one of the continuous-flow systems and then sifted it through 0.5cm wire mesh. We created worm juice for the experiment each week using extracts from household operations. To do this, we identified ∼12 households that had well-established composting systems (worms were thriving, food waste was being consistently added) that were protected from the elements (to prevent rainwater from entering the system). We collected the liquid that collected in the bottom bin of each two-compartment system and then combined all collected extracts in a 25L container. We then filtered and diluted this liquid (9 water:1 extract) before adding it to plots. Water used in this dilution came from a local borehole.

### Site preparation and experimental design

We established 54 square production plots on the central site in June 2025 (Figure 1D). We made all plots 0.25m^2^ and arranged them in a grid with each plot separated by 50cm. We first cleared all vegetation (mostly grasses) and sifted soil in each plot to 30cm depth to remove rocks and debris. Soil was sandy (see below), as is typical in Khayelitsha. Soil conditions appeared to vary slightly across the plot, so we organized plots into 9 blocks based on location within the site.

Within each block, we created a fully factorial design with three levels of vermicompost and two levels of worm juice. We allocated plots equally to one of three vermicompost treatments: 0% (control), 10% (by volume) vermicompost, and 20% vermicompost. We created vermicompost treatments by mixing all soil excavated from a plot with finished vermicompost from the continuous-flow systems. In addition, within each vermicompost treatment, we randomly allocated plots to either receive or not receive (“controls”) weekly worm juice supplements. Three weeks after initial planting of seedlings, we began adding 2L worm juice per week to each treatment plot, and 2L of water per week to control plots. We did not add any other water to plots.

We initiated plots in mid-June 2025. We planted green Swiss chard (*Beta vulgaris* sub-species *vulgaris*) from seedlings purchased from a local distributor. This plant is probably the most common winter crop grown in Khayelitsha. We planted seedlings in small depressions after placing a small amount of vermicompost at the bottom of each depression (following methods used by local farmers). We added four seedlings per plot in a grid formation, each separated by 15cm (and at least 5cm from plot edges). We then mulched all spaces between plots using sawdust (Figure 1D). We did not add mulch within plots, following practices used by local farmers. We also planted ∼100 seedlings in a separate, unfertilized plot to serve as replacement plants if any seedlings in the experiment did not establish.

Within the first 10 days of the experiment, we replaced 12 of 216 of the initial seedlings with these replacements. No plants failed after the initial 10-day period.

Three weeks after initiating plots, we collected 3 samples each of control soil, 10% vermicompost soil, 20% vermicompost soil, and pure vermicompost. We sent all samples to the Soil, Water, and Plant laboratory of the Western Cape Department of Agriculture. Soils were analyzed for pH(KCl), resistance, texture, calcium, magnesium, potassium, sodium, citric acid concentration, and total cations.

We harvested chard plants 72 days after planting. We cut plants at the base and determined wet mass on a digital scale (±1g).

We conducted all statistical analyses using the lme and emmeans packages in R (version 4.5.2). We used total plot wet mass in the statistical analysis of yield. We used linear mixed-effects models to account for the block design. We treated vermicompost amendment level (0%, 10%, 20%), worm juice application (present or absent), and their interaction as fixed effects and included block as a random intercept. We assessed significance using ANOVA and evaluated model assumptions using visual inspection of residual plots. Pairwise comparisons were evaluated with Tukey-adjusted contrasts using the emmeans package. We evaluated differences in soil pH among vermicompost treatments using a one-way ANOVA, with vermicompost percentage treated as a fixed factor, followed by pairwise comparisons when ANOVAs were significant. We tested for normality of residuals using the Shapiro– Wilk test, which indicated no deviation from normality for any soil metric.

## Results

Amendments with vermicompost and worm juice created in a community composting project both significantly increased Swiss chard yield (Figure 3). The vermicompost effect was particularly strong (F_2,40_ = 14.32, P < 0.0001), as yields in the 10% and 20% vermicompost treatments were on average 51.2% and 58.5% higher, respectively, than yields in negative control plots. The effect of worm juice addition was also significant (F_1,40_ = 5.52, P = 0.024), but less substantial (18.3% yield increase). The vermicompost-by-worm juice interaction (F_2,40_ = 2.34, P = 0.110) was not significant.

**Figure 3.**
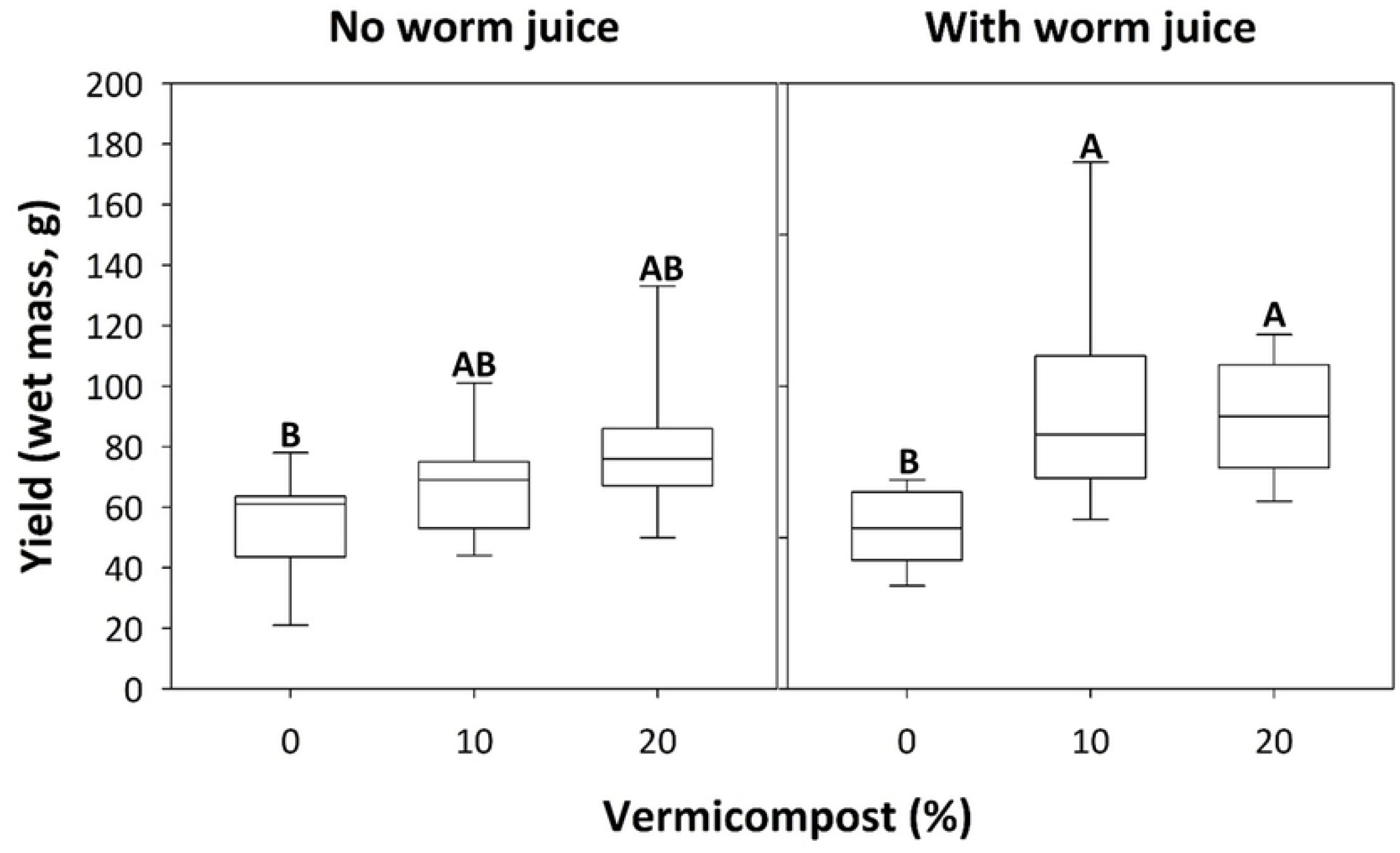
Yield results (wet mass, in grams) from the growing experiment. Swiss chard was grown in either unamended soil (0% vermicompost), with 10% (by volume) vermicompost, or with 20% vermicompost added to plots before planting. For each vermicompost treatment, half of plots either did not or did receive weekly additions of worm juice. Letters above bars indicate significant (p < 0.05) differences in pairwise comparisons.

Amendments with vermicompost affected several soil properties (Table 1). Unamended soil at this location was sandy, had high pH, high resistance, and moderately low levels of calcium, magnesium, potassium, sodium, phosphorus, and total cations. Finished vermicompost had significantly lower pH (overall ANOVA - F_3,8_ = 46.97, P < 0.0001) and resistance (F_3,8_ = 91.3, P < 0.0001), and significantly higher levels of total cations (F_3,8_ = 257.2, P < 0.0001) and all measured nutrients (Ca (F_3,8_ = 210, P < 0.0001), Mg (F_3,8_ = 1350, P < 0.0001), K (F_3,8_ = 142.6, P < 0.0001), Na (F_3,8_ = 61.93, P < 0.0001), P (F_3,8_ = 1398, P < 0.0001)). The 10% and 20% vermicompost soils also had significantly lower pH and resistance compared to the unamended soil, and higher levels of P (Table 1). Amendments did not significantly affect other soil metrics, although statistical comparisons are based on only 3 samples per treatment.

**Table 1.** Results (mean ± 1 S.E.) from analyses of soil samples collected from experimental plots and vermicompost from the continuous-flow system three weeks after the establishment of the growing experiment. Letter superscripts indicate significant (p < 0.05) differences in pairwise comparisons.

| Soil feature | units | % vermicompost |  |  |  |
| --- | --- | --- | --- | --- | --- |
|  |  | 0 | 10 | 20 | 100 |
| **pH(KCl)** |  | 8.17±0.08<sup>a</sup> | 7.90±0.06<sup>b</sup> | 7.67±0.03<sup>b</sup> | 7.23±0.03<sup>c</sup> |
| **Resistance** | ohms | 2,116.7±62<sup>a</sup> | 1,276.7±103<sup>b</sup> | 1,070.0±120<sup>c</sup> | 126.7±3.3<sup>d</sup> |
| **Texture** |  | sand | sand | sand | compost |
| **Calcium** | cmol(+)/kg | 17.5±0.31<sup>a</sup> | 18.1±0.25<sup>a</sup> | 18.0±0.30<sup>a</sup> | 29.3±0.60<sup>b</sup> |
| **Magnesium** | cmol(+)/kg | 3.5±0.01<sup>a</sup> | 3.8±0.11<sup>a</sup> | 4.1±0.20<sup>a</sup> | 14.8±0.19<sup>b</sup> |
| **Potassium** | mg/kg | 47.7±7.3<sup>a</sup> | 72.7±8.3<sup>a</sup> | 135.0±26<sup>a</sup> | 5299.3±435<sup>b</sup> |
| **Sodium** | mg/kg | 193.3±1.45<sup>a</sup> | 188.0±2.31<sup>a</sup> | 190.0±1.0<sup>a</sup> | 962.3±98<sup>b</sup> |
| **P (citric acid)** | mg/kg | 195.0±33<sup>a</sup> | 240.7±21<sup>b</sup> | 328.7±29<sup>bc</sup> | 2133.0±19<sup>c</sup> |
| **Total cations** | cmol(+)/kg | 22.0±0.08<sup>a</sup> | 23.0±0.08<sup>a</sup> | 23.3±0.08<sup>a</sup> | 61.9±0.08<sup>b</sup> |

## Discussion

We demonstrated that vermicompost and worm juice generated from food waste in a community composting project in a South African township increased Swiss chard yield by 50-60%. This result suggests that low-tech, low-cost projects can help meet pressing needs of urban farmers for local sources of materials for soil enrichment.

Vermicompost addition appeared to partially offset deficiencies in the parent soil (Table 1). Unamended soil pH (∼8.2) was far more alkaline than recommended soil pH (6.0-7.0) for chard production (Knežević et al. 2014). Addition of vermicompost significantly lowered the pH is treatment plots (7.67 and 7.9), but further amendments are needed to reduce pH to optimal conditions for this crop. Unamended soil also had levels of key soil elements (calcium, magnesium, potassium, sodium, and phosphorus) and total cations that were considerably lower than recommended levels for Swiss chard production (Kebede et al. 2023). Vermicompost addition significantly increased soil phosphorus levels but did not affect other soil measured soil parameters, suggesting that significant leaching occurred in this sandy soil. These results suggest that 1) even moderate soil property changes caused by vermicompost addition can substantially increase yields in this system, and 2) there are ample opportunities for additional soil amendment to further increase production. One promising opportunity is to mix biochar with vermicompost to increase water retention and reduce nutrient leaching (Ehsanul et al. 2023).

These results are somewhat preliminary, and several caveats are in order when interpreting their significance. First, the production of the vermicompost used in the study cannot be exactly replicated because inputs into the household and central-site composting systems were not tightly regulated. We gave households instructions about best practices for vermicomposting, including avoidance of adding citrus, onion, and meat scraps into bins. Although weekly visits suggested that households followed these instructions without fail, we have no information about the quantity and mixture of fruit and vegetable waste added to bins, which in turn produced the pre-compost added to our continuous-flow system. Second, we have no information about parent soil characteristics beyond data presented in Table 1, and other factors such as various soil toxins may interact with vermicomposting products to affect yields. Finally, our results are based on yields for one crop during one growing season at one location, with application of the one set of growing practice used by local farmers (e.g., no mulch added, no chemical application for pest control). Nevertheless, the >50% yield increase with vermicompost and worm juice application suggests that such a grassroots, community-embedded composting operation can increase production and nutritional gains for urban farmers in this community. Future work will explore how additional low-cost inputs can be combined with vermicompost and worm juice in this system to increase yields for urban farmers.

Scaling up this and similar community-embedded vermicomposting operations may provide multiple benefits to marginalized areas of SSA cities and beyond. Discovering and implementing other grassroots, low-cost, community-embedded approaches for increasing yields and net returns from community and home gardens can encourage more and broader participation in urban farming, which has been shown to provide multiple social and environmental benefits in poor communities (Kanosvamhira 2024, Kanosvamhira et al. 2025). More work is needed, however, to optimize these low-tech approaches to maximize gains for practitioners and to facilitate their adoption in other communities.

## Acknowledgements

This work was funded by National Science Foundation Undergraduate Biology Education, Grant/ Award Number DEB-2018837 and by the University of St. Thomas (MN)

## References

1. Büttner N, Stalder S, Volpi M, Suel E, Harttgen K. Large-scale slum mapping in sub-Saharan Africa’s major cities: Remote sensing and deep learning reveal strong slum growth in the urban periphery between 2016 and 2022. Habitat International. 2025;161:103403.

2. Cerda A, Artola A, Font X, Barrena R, Gea T, Sánchez A. Composting of food wastes: Status and challenges. Bioresour Technol. 2018;248A:57–67.

3. Chalak A, Abou-Daher C, Chaaban J, et al. The global economic and regulatory determinants of household food waste generation: A cross-country analysis. Waste Manag. 2016;48:418–422.

4. Ehsanul K, Kim KH, Eilhann EK. Biochar as a tool for the improvement of soil and environment. Front Environ Sci. 2023;11:1324533.

5. Elmqvist T, Setälä H, Handel SN, van der Ploeg S, Aronson J, Blignaut JN, et al. Benefits of restoring ecosystem services in urban areas. Curr Opin Environ Sustain. 2015;14:101–108.

6. Fan G, Yu Y, Gao M, Sharifi A, Pathak M, Zhang K, et al. One climate, one urban, one health. Sustain Cities Soc. 2025;130:106619.

7. G VB, Sonba MS. Livelihoods for urban slums and economic growth. Afr J Biomed Res. 2024;27:2570– 2580.

8. Ganapathy NRV, Elango AC, Balaji G, Sankaranarayanan M, Sharma M. A comprehensive review of earthworm-derived vermiproducts and their role in sustainable agriculture. Discov Appl Sci. 2025;7:995.

9. Garg VK, Suthar S, Yadav A. Management of food industry waste employing vermicomposting technology. Bioresour Technol. 2012;126:437–443.

10. Kanosvamhira TP. How do we get the community gardening? Grassroots perspectives from urban gardeners in Cape Town, South Africa. J Cult Geogr. 2023;40:47–63.

11. Kanosvamhira TP. Exploring urban community gardens as ‘third places’: fostering social interaction in distressed neighbourhoods of Cape Town, South Africa. Leis Stud. 2024;43:1–18.

12. Kanosvamhira TP, Shade MD. Urban agriculture for environmentally just cities: The case of urban community gardens in Cape Town, South Africa. Local Environ. 2024;30:133–149.

13. Kanosvamhira TP, Musasa T, Mupepi O. The potential for urban agriculture in Cape Town, South Africa: A suitability analysis. Ann GIS. 2025;31:107–122.

14. Kebede T, Diriba D, Boki A. The effect of organic solid waste compost on soil properties, growth, and yield of Swiss chard (Beta vulgaris L.). Sci World J. 2023;2023:6175746.

15. Kiribou R, Bedadi B, Dimobe K, Ndemere J, Neya T, Ouedraogo V, Dejene SW. Urban farming system and food security in sub-Saharan Africa: Analysis of the current status and challenges. Urban Agric Reg Food Syst. 2024;9:e70007.

16. Knežević M, Djurovic D, Mugoša B, Strunjaš M, Topalovic A. Relationships between parameters of soil and chard (Beta vulgaris L. var. cicla). Agric For. 2014;60:275–283.

17. Kronenberg J, Andersson E, Elmqvist T, Łaszkiewicz E, Xue J, Khmara Y. Cities, planetary boundaries, and degrowth. Lancet Planet Health. 2024;8:e234–e241.

18. Lynge H, Visagie J, Scheba A, Turok I, Everatt D, Abrahams C. Developing neighbourhood typologies and understanding urban inequality: A data-driven approach. Reg Stud Reg Sci. 2022;9:618–640.

19. Marx B, Stoker T, Suri T. The economics of slums in the developing world. J Econ Perspect. 2013;27:187– 210.

20. McPhearson T, Pickett STA, Grimm NB, Niemelä J, Alberti M, Elmqvist T, et al. Advancing urban ecology toward a science of cities. BioScience. 2016;66:198–212.

21. Mmereki D, Baldwin A, Li B. A comparative analysis of solid waste management in developed, developing and lesser developed countries. Environ Technol Rev. 2016;5:120–141.

22. Mmereki D, David VE Jr, Brownell AHW. The management and prevention of food losses and waste in low- and middle-income countries: A mini-review in the Africa region. Waste Manag Res. 2024;42:287– 307.

23. Modibedi TP, Masekoameng MR, Maake MM. The contribution of urban community gardens to food availability in Emfuleni Local Municipality, Gauteng Province. Urban Ecosyst. 2021;24:301–309.

24. Musosa L, Shekede MD, Gwitira I, Chirisa I, Tevera D, Matamanda AR. Auditing the spatial and temporal changes in urban cropland in Harare Metropolitan Province, Zimbabwe. Afr Geogr Rev. 2024;43:170– 185.

25. Mwakiwa E, Maparara T, Tatsvarei S, Muzamhindo N. Is community management of resources by urban households feasible? Lessons from community gardens in Gweru, Zimbabwe. Urban For Urban Green. 2018;34:97–104.

26. Olivier DW. Urban agriculture promotes sustainable livelihoods in Cape Town. Dev South Afr. 2019;36:17–32.

27. Potts D. Debates about African urbanisation, migration and economic growth: What can we learn from Zimbabwe and Zambia? Geogr J. 2016;182:251–264.

28. Reuther S, Dewar N. Competition for the use of public open space in low-income urban areas: The economic potential of urban gardening in Khayelitsha, Cape Town. Dev South Afr. 2006;23:97–122.

29. Rigon A. Diversity, justice and slum upgrading: An intersectional approach to urban development. Habitat Int. 2022;130:102691.

30. Schröder C, Häfner F, Larsen OC, Krause A. Urban organic waste for urban farming: Growing lettuce using vermicompost and thermophilic compost. Agronomy. 2021;11:1175.

31. City of Cape Town. 2011 Census: Khayelitsha Health District. Cape Town: City of Cape Town; 2013.

32. Sheahan M, Barrett CB. Food loss and waste in Sub-Saharan Africa: A critical review. Food Policy. 2017;70:1–12.

33. Sherman R. The Worm Farmer’s Handbook: Mid-to Large-Scale Vermicomposting for Farms, Businesses, Municipalities, Schools, and Institutions. White River Junction (VT): Chelsea Green Publishing; 2018.

34. Swanepoel JW, Van Niekerk JA, Tirivanhu P. Analysing the contribution of urban agriculture towards urban household food security in informal settlement areas. Dev South Afr. 2021;38:785–798.

35. Wang F, Zhang Y, Su Y, Wu D, Xie B. Pollutant control and nutrient recovery of organic solid waste by earthworms: Mechanism and agricultural benefits of vermicomposting. J Environ Chem Eng. 2024;12:112610.

36. Webb D. “These aren’t the jobs we want”: Youth unemployment and anti-work politics in Khayelitsha, Cape Town. Soc Dyn. 2021;47:1–17.

37. Williams DS, Costa MM, Sutherland C, Celliers L, Scheffran J. Vulnerability of informal settlements in the context of rapid urbanization and climate change. Environ Urban. 2019;31:157–176.

38. Zimmer A, Guido Z, Tuholske C, Pakalniskis A, Lopus S, Caylor K, Evans T. Dynamics of population growth in secondary cities across southern Africa. Landscape Ecol. 2020;35:2501–2516.

